# Physical phenotype of blood cells is altered in COVID-19

**DOI:** 10.1101/2021.02.12.429482

**Authors:** Markéta Kubánková, Bettina Hohberger, Jakob Hoffmanns, Julia Fürst, Martin Herrmann, Jochen Guck, Martin Kräter

## Abstract

Clinical syndrome coronavirus disease 2019 (COVID-19) induced by severe acute respiratory syndrome coronavirus 2 (SARS-CoV-2) is characterized by rapid spreading and high mortality worldwide. While the pathology is not yet fully understood, hyper-inflammatory response and coagulation disorders leading to congestions of microvessels are considered to be key drivers of the still increasing death toll. Until now, physical changes of blood cells have not been considered to play a role in COVID-19 related vascular occlusion and organ damage. Here we report an evaluation of multiple physical parameters including the mechanical features of five frequent blood cell types, namely erythrocytes, lymphocytes, monocytes, neutrophils, and eosinophils. More than 4 million blood cells of 17 COVID-19 patients at different levels of severity, 24 volunteers free from infectious or inflammatory diseases, and 14 recovered COVID-19 patients were analyzed. We found significant changes in erythrocyte deformability, lymphocyte stiffness, monocyte size, and neutrophil size and deformability. While some of these changes recovered to normal values after hospitalization, others persisted for months after hospital discharge, evidencing the long-term imprint of COVID-19 on the body.

## Introduction

Peripheral blood is a key body fluid analyzed during the diagnostic routine, including infectious disease diagnostics. Infection by severe acute respiratory syndrome coronavirus 2 (SARS-CoV-2) may lead to the clinical syndrome coronavirus disease 2019 (COVID-19), which is accompanied by changes in numbers and phenotypes of blood cells (1). Typically, various immune cells such as T-lymphocytes, monocytes and macrophages get activated and contribute to the so-called hyper-inflammatory response (2). The uncontrolled inflammation is believed to be a major cause of disease severity and death during COVID-19 (3). Furthermore, abnormal coagulation and thrombotic events leading to vascular occlusion are described as major contributors to the high mortality (4, 5).

Apart from the biochemical state of blood cells, infectious diseases can also alter their physical properties, including morphological or mechanical features. It is long known that mechanical properties of cells can act as a disease marker, as reviewed by Di Carlo (6), and can contribute to vascular occlusion, as reviewed by Lipowsky (7). To date, a systematic evaluation of the effect of COVID-19 on the physical phenotypes of the most frequent blood cells was missing. To address this gap, we employed real-time deformability cytometry (RT-DC), a label free, high-throughput technology that allows quick image-based mechanical interrogation of cells at rates of up to 1000 cells per second (8). Previously, the technique was used to detect disease specific signatures of blood cell pathological changes in several conditions including spherocytosis, malaria, acute lymphoid leukemia, and acute myeloid leukemia (9). During viral respiratory tract infection Toepfner *et* al. reported an increase of neutrophil and monocyte size and deformability, as well as larger and more deformable lymphocytes in acute Epstein-Barr-virus infection.

Here, we examine COVID-19 related changes of physical phenotype of several peripheral blood cell types, namely erythrocytes, lymphocytes, monocytes and neutrophils. In total, more than 4×10^6^ blood cells from 55 blood samples were analyzed, including 17 COVID-19 positive patients, 14 recovered patients, and 24 age matched volunteers showing no indication of infection or inflammatory disease. We found that COVID-19 is linked with significantly decreased lymphocyte stiffness, increased monocyte cell size, the appearance of smaller and less deformable erythrocytes, and the presence of large, deformable, activated neutrophils. Certain changes had not returned to the control group levels months after release from the hospital, bringing evidence of the long-lasting effects of COVID-19 on the circulatory system. Our results show that real-time deformability cytometry can be used to follow the course of COVID-19 and the immune response against it. In the future, we anticipate that measurements of morphological and mechanical properties of blood cells will contribute towards improving infectious disease diagnostics.

## Results and Discussion

We studied the peripheral blood from 17 COVID-19 patients hospitalized at the time of sample collection (median age 68 ± 10.4 years) compared to a cohort of 24 volunteers free from infectious or inflammatory diseases (62.5 ± 13.6 years) and 14 blood donors on average 7 months after hospitalization with COVID-19 (age 58.6 ± 12.4 years, herein referred to as the ‘recovered’ patients). Whole blood was diluted in measurement buffer at a ratio of 1:20 and analyzed with RT-DC (Figure 1). For each patient, the five most frequent blood cell populations were manually gated according to the established analysis protocol (9): erythrocytes, lymphocytes, monocytes, neutrophils, and eosinophils.

**Figure 1.**
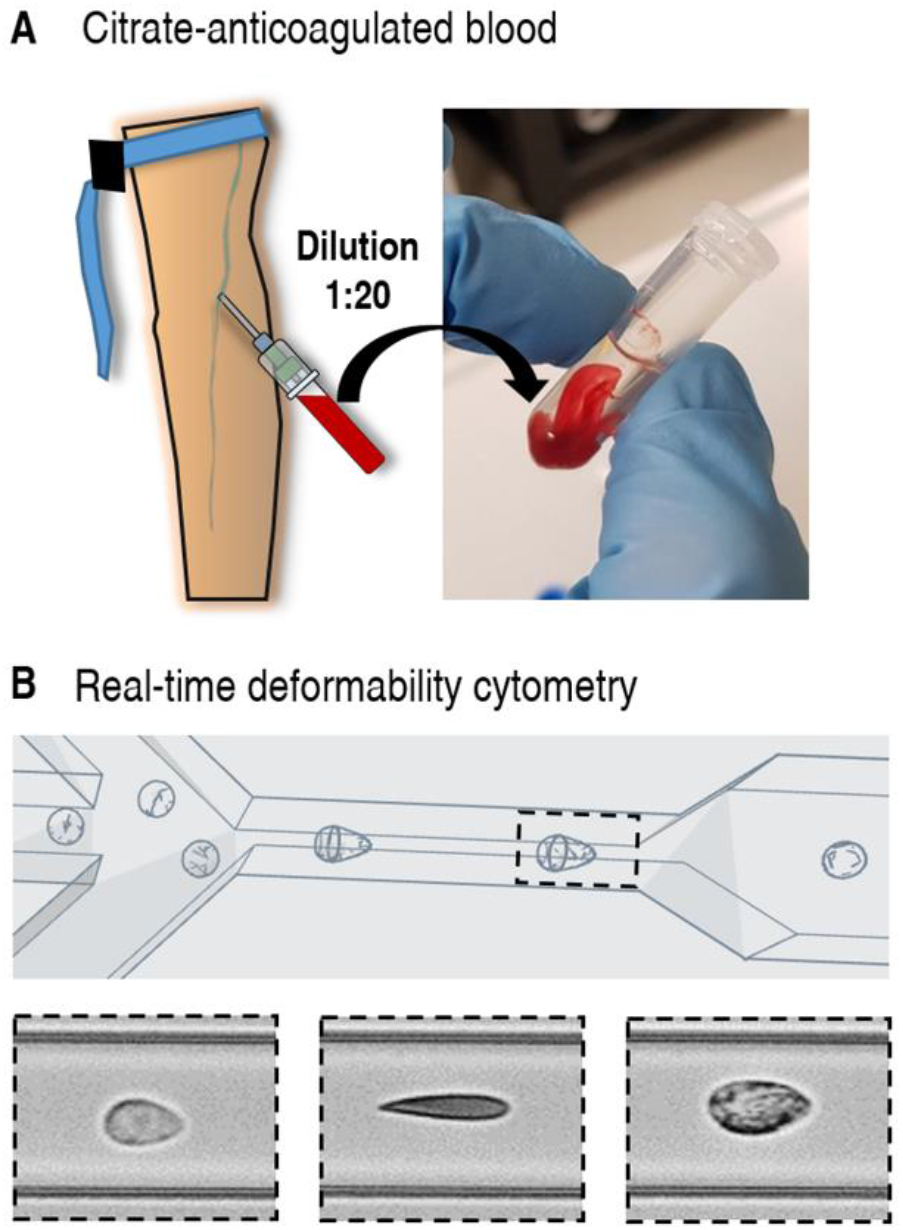
Scheme of an RT-DC measurement of a peripheral blood sample. A) Venous citrate-anticoagulated blood was obtained, of which 50 µl was diluted in 950 µl of measurement buffer consisting of PBS and methyl cellulose. The tube was shaken gently until the mixture appeared homogenous. B) The blood cell suspension was pumped through a microfluidic chip mounted on an inverted microscope and images were processed in real-time to obtain the physical parameters of each cell.

In agreement with other studies (10, 11) we observed significant alterations of white blood cell (WBC) counts in severe COVID-19 cases, namely neutrophilia (elevated number of neutrophils) and lymphopenia (decreased number of lymphocytes), Supplementary figure 1. The median neutrophil to lymphocyte ratio (NLR) increased from 0.97 ± 0.70 to 3.62 ± 3.36, with several cases where NLR was over 8. The NLR increase calculated from RT-DC data was consistent with literature, where NLR is reported as a prognostic marker of COVID-19 mortality (12). In some patients we also observed monocytosis (13): while the proportion of monocytes to total WBC was mostly in the normal range of 2–8%, in several COVID-19 cases it reached over 10% (Supplementary figure 1). No significant changes were found in eosinophil counts. Overall these findings confirm that, purely from the images of cells obtained with RT-DC, it is possible to reproduce the results of conventional complete blood counts (9). The focus of this study, however, was the interrogation of changes of blood cell physical phenotype. The following sections describe alterations of erythrocytes and leukocytes during COVID-19.

### COVID-19 induces the appearance of erythrocytes with distinct physical phenotype

RT-DC analysis revealed erythrocyte anomaly in COVID-19 patients, mainly characterized by the appearance of erythrocytes with low deformation in standardized channel flow conditions (Figure 2 A-E). The median deformation of erythrocytes exhibited a weak decrease in COVID-19 patients compared to healthy donors and recovered patients (Kruskal-Wallis *p* = .22, *χ*^2^= 3.1, ε^2^= 0.06), Figure 2 F. It is noteworthy that several of the COVID-19 patients had very low median erythrocyte deformability compared to the rest of the blood donors.

**Figure 2.**
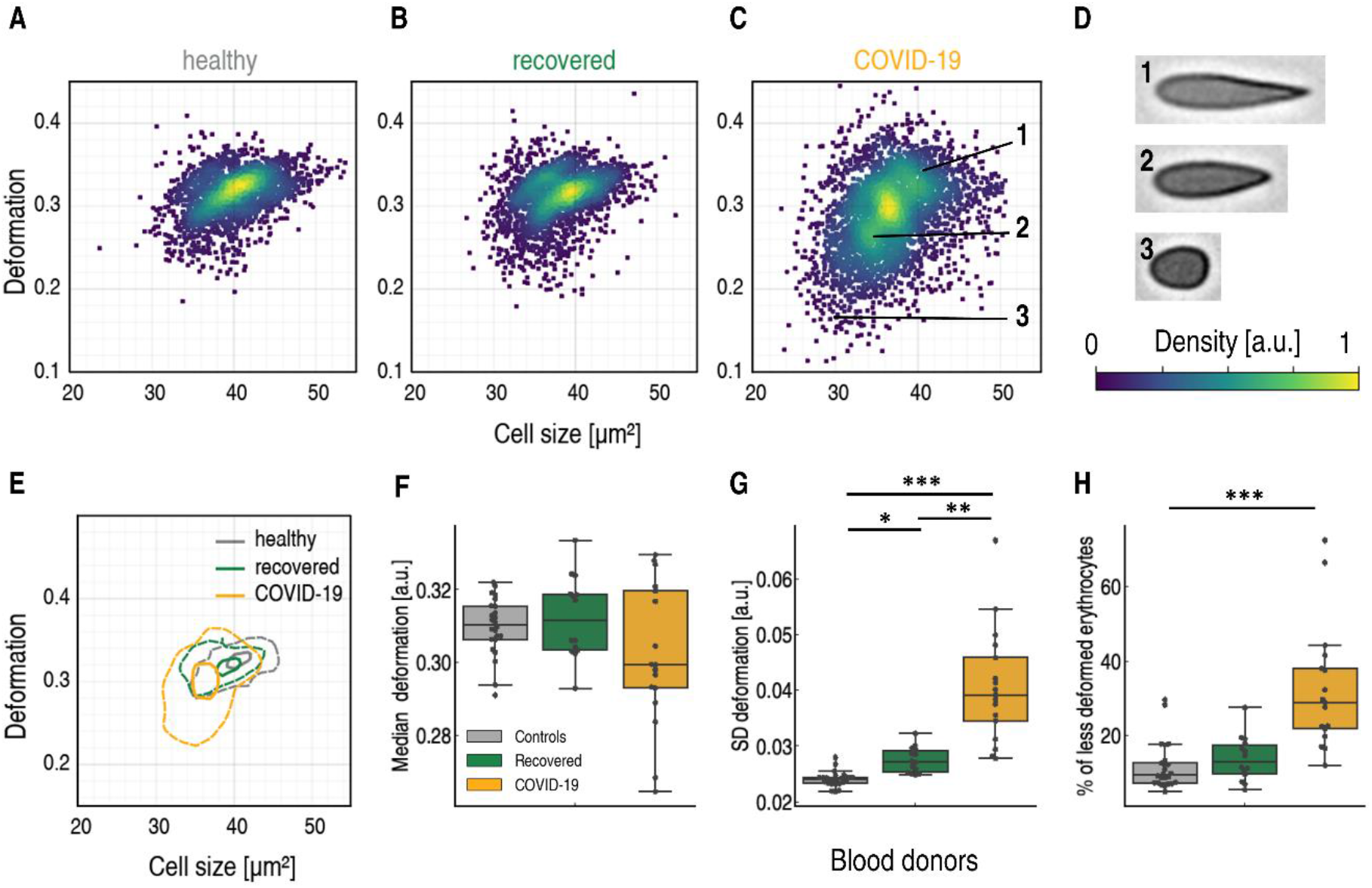
Decreased size and deformability of erythrocytes of hospitalized COVID-19 patients. Typical scatter plot of erythrocyte deformation vs. cell size (cross-sectional area) of a healthy blood donor with no known viral infection (A) compared to a patient four months after undergoing COVID-19 (B) and a patient with COVID-19 in an intensive care unit (C). The erythrocytes shown in (D) are representative images of cells in the clusters marked by corresponding numbers in the scatter plot. (E) Kernel density estimate plots demonstrating the differences in cell size and deformability between the three donors (A-C). The comparison of median values of deformation (F) and standard deviation of deformation (G) between the control group of blood donors (n = 24), recovered patients (n = 14), and patients hospitalized with COVID-19 (n = 17). (H) The percentage of erythrocytes with deformation below an arbitrary threshold of 0.28. The fraction of these less deformable erythrocytes in the total erythrocyte count is significantly increased in COVID-19 patients. Statistical comparisons were done using Kruskal-Wallis test with Dunn’s posthoc test, * *p* < .05, ** *p* < .01, *** *p* < .001.

Significant differences with strong effect size were observed in the standard deviations of erythrocyte deformation (Kruskal-Wallis *p* < .0001, *χ*^2^ = 42.3, ε^2^ = 0.78), Figure 2 G. The significant broadening of the deformation distribution during COVID-19 was the result of the appearance of erythrocytes with low deformability, as shown in Figure 2 D. Such cells were very rare in the healthy and recovered patient cohorts. The percentage of erythrocytes with deformation under 0.28 was on average 10.6% for healthy donors, 14.6% for recovered patients, and a striking 32.4% for hospitalized patients (*p* < .0001, *χ*^2^ = 27.0, ε^2^ = 0.53), Figure 2 H.

Alongside the significant difference in standard deviation (SD) of deformation between hospitalized patients and the healthy cohort (*p* < .0001, see Supplementary table 2 for detailed results of Dunn’s post-hoc tests), we also found significant differences between recovered and hospitalized cohorts (*p* = .002) and between recovered and healthy cohorts (*p* = .03)., Figure 2 G. Clearly, the erythrocytes of the recovered patients had not fully returned to the state of the healthy cohort.

In addition to the increased SD of deformation, we also found increased SD of cell size, specifically the cross-sectional area of the image-derived cell contours, and the cell volume (Supplementary figure 2). The increased standard deviations of cell size and deformation were in accordance with reported broadening of the red blood cell distribution width (RDW), a routine complete blood count component (14).

In general, the phenotype changes observed with RT-DC may be associated with structural and functional changes. A proteomics study has reported that COVID-19 causes irreversible damage to the erythrocyte proteome (15). The authors found that oxidative stress connected with COVID-19 damages essential proteins in erythrocytes, including those that influence membrane structure and the ability to transport and deliver oxygen. Since mature erythrocytes cannot synthesize new proteins to replace damaged ones, and the average lifespan of erythrocytes is 120 days, the authors hypothesize that the circulation of irreversibly damaged erythrocytes with impaired function could contribute to the long-term effects of COVID-19 (15).

According to our findings, COVID-19 is connected with the appearance of less deformable erythrocytes. Cell deformability is thought to be a key factor determining splenic clearance (16) and it is likely that erythrocytes strongly deviating from normal deformability get removed by the spleen.

However, we observed that erythrocyte deformability in recovered patients had not returned to healthy donor levels. Thus, we hypothesize that erythrocytes with only minor deviation from normal deformability pass through the spleen unnoticed. Due do the long erythrocyte lifespan, such cells may remain in the blood circulation for months, possibly contributing to the problems experienced by post-COVID long haulers (17).

### Decreased lymphocyte stiffness in COVID-19 patients

RT-DC analysis revealed high deformability of peripheral blood lymphocytes during severe COVID-19 (*p* = .013, *χ*^2^ = 8.7, with relatively strong effect size ε^2^ = 0.16), as can be seen in Figure 3 A-E. While lymphocyte cell size did not differ among healthy donors, recovered, and hospitalized COVID-19 patients (medians 37.8 ± 1.3 µm^2^, 38.6 ± 0.7 µm^2^ and 39.0 ± 2 µm^2^, respectively, Figure 3 F), lymphocyte deformation in standardized channel flow conditions was elevated during COVID-19. Compared to the healthy donor median deformation of 0.025 ± 0.006, hospitalized COVID-19 patient lymphocytes had a significantly higher median deformation 0.029 ± 0.003, *p* = .011 (Figure 3 G). Lymphocyte deformation was 0.026 ± 0.002 for recovered patients, not significantly different from that of the control group.

**Figure 3.**
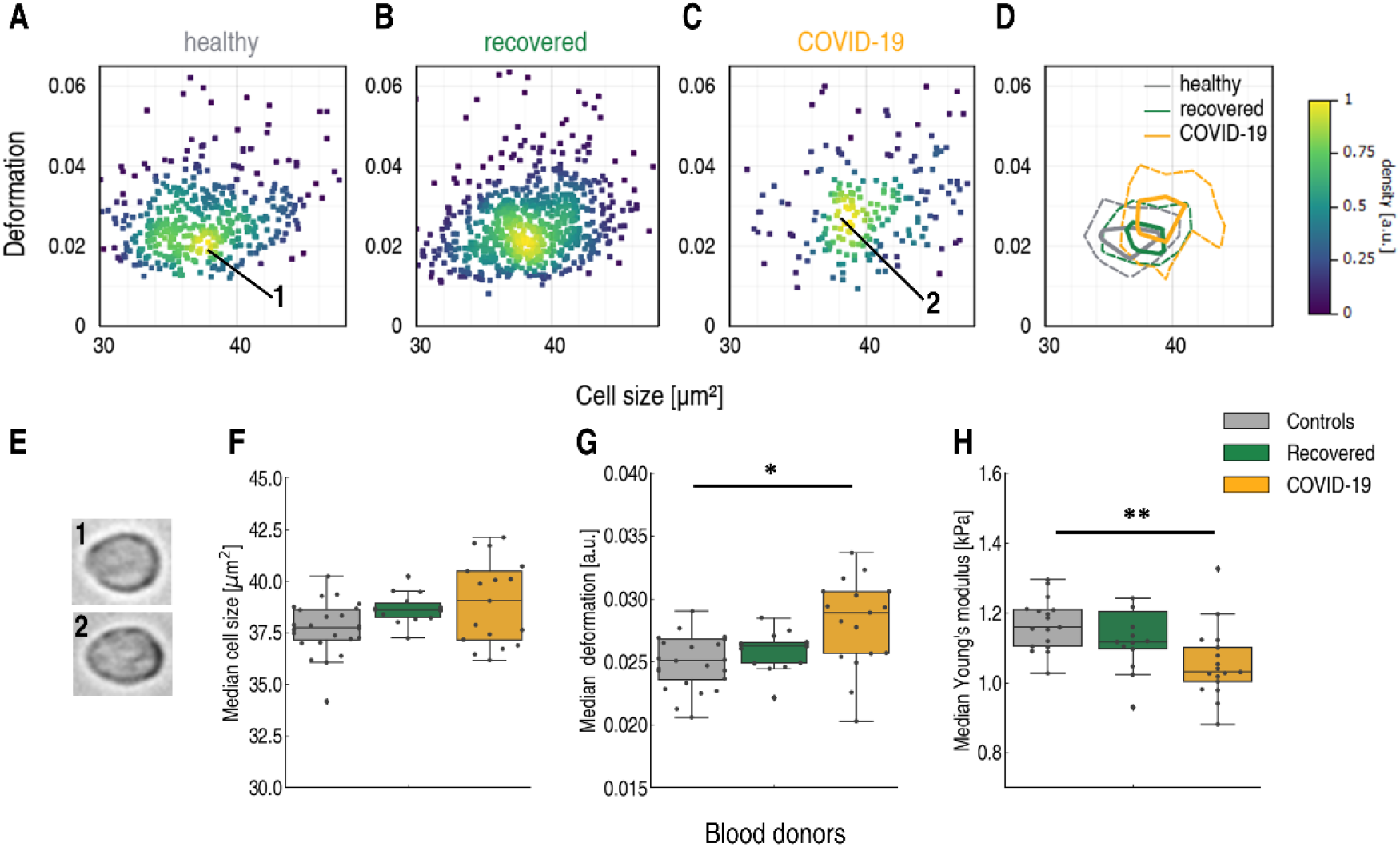
Lymphocytes are less stiff in peripheral blood of hospitalized COVID-19 patients. Typical scatter plot of lymphocyte deformation vs. cell size (cross-sectional area) of a a healthy blood donor with no known viral infection (A) compared to a patient four months after undergoing COVID-19 (B) and a patient with COVID-19 in an intensive care unit (C). (D) Kernel density estimate plots demonstrating the differences in cell size and deformation among the three donors (A-C). (E) No significant differences in lymphocyte cell size were found between healthy blood donors (grey, n = 24), recovered patients approximately five months after undergoing COVID-19 (green, n = 14), and patients hospitalized with COVID-19 (yellow, n = 17). (F) Lymphocytes exhibit significantly increased deformation in hospitalized COVID-19 patients. (G) Young’s modulus of lymphocytes is significantly higher in COVID-19 patients compared to the healthy or recovered donors. Statistical comparisons were done using Kruskal-Wallis test with Dunn’s posthoc test, * *p* < .05, ** *p* < .01, *** *p* < .001.

The sphericity of lymphocytes under normal conditions allowed us to exploit the developed theoretical framework (18) to calculate the Young’s modulus from RT-DC data. The Young’s modulus, a measure of overall cell stiffness, was significantly lower in the COVID-19 cohort (*p* = .003, *χ*^2^ = 11.7, ε^2^ = 0.22) (Figure 3 H). While the control group median Young’s modulus was 1.15 ± 0.12 kPa, it went down to 1.03 ± 0.10 kPa in hospitalized COVID-19 patients (*p* = .003, Dunn’s post-hoc test). To the best of our knowledge, this study reveals the first evidence of altered mechanical properties of lymphocytes during COVID-19.

### Monocytes of COVID-19 patients exhibit a dramatic increase in cell volume

In inflammatory disease, monocytes can contribute to the immune response either directly or via differentiation into dendritic cells or macrophages. Therefore it is not surprising that altered monocyte phenotype and function is characteristic for COVID-19 patients (1). Examination of monocytes with RT-DC revealed a significant change in monocyte size (*p* < .0001, *χ*^2^ = 30.6, ε^2^ = 0.57) triggered by the appearance of larger monocytes during COVID-19 (Figure 4 A-D). Monocytes of hospitalized COVID-19 patients had a median cell cross-sectional area of 70.5 ± 7.1 µm^2^, significantly higher compared to recovered patients (65.0 ± 2.5 µm^2^, *p* < .0001, Dunn’s post-hoc test) and that of the healthy cohort, 63.8 ± 2.2 µm^2^ (*p* < .0001), Figure 4 E. This represents a 9.5% increase from the median cross-sectional area of the healthy cohort. In addition, the standard deviation of cross-sectional area increased during COVID-19 (*p* = .001, *χ*^2^ = 13.7, ε^2^ = 0.25) due to the appearance of large monocytes (Figure 4 F).

**Figure 4.**
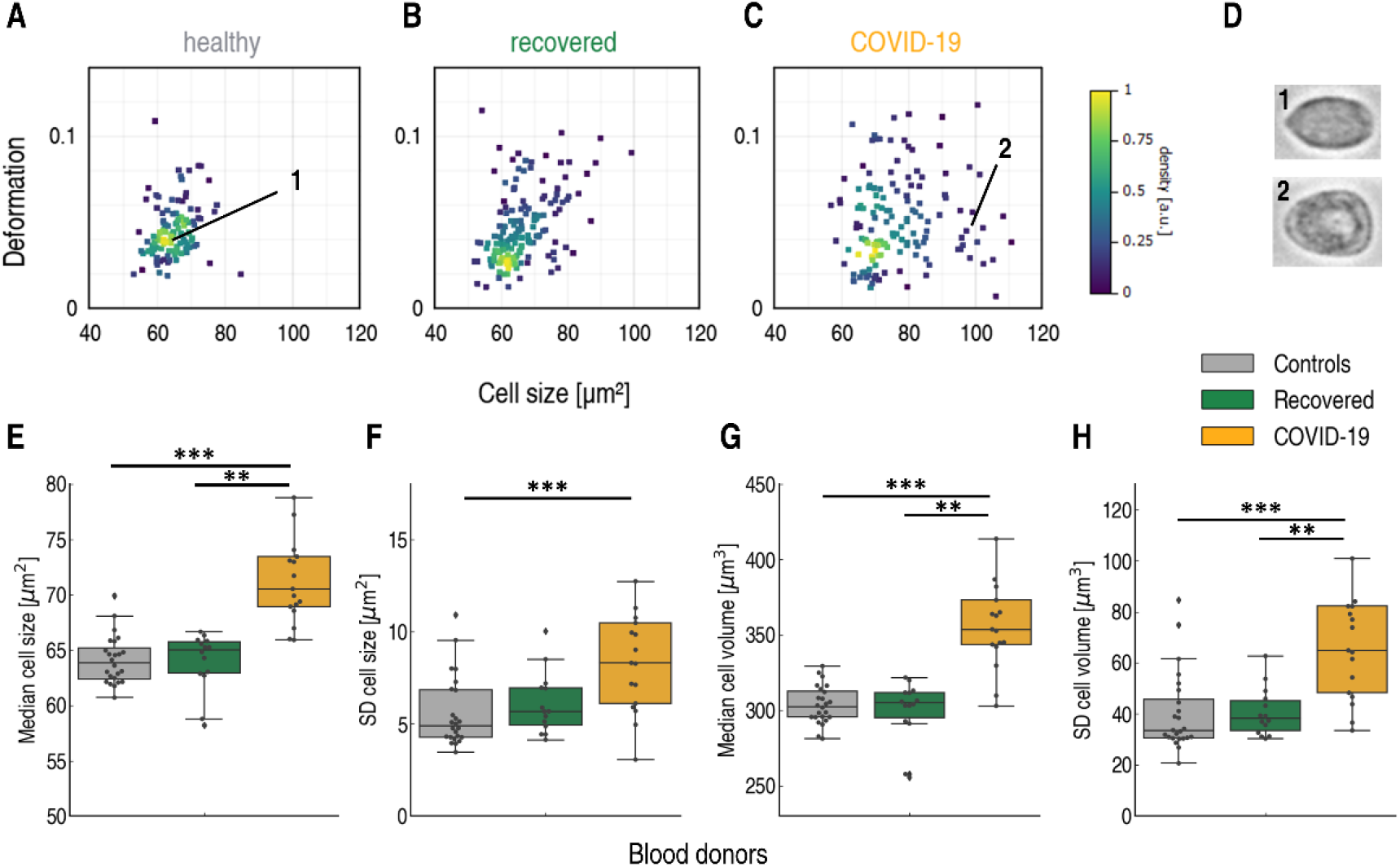
The appearance of large monocytes in COVID-19 patients. Typical scatter plot of monocyte deformation vs. cell size (cross-sectional area) of a healthy blood donor with no known viral infection (A) compared to a patient four months after undergoing COVID-19 (B) and a patient with COVID-19 in an intensive care unit (C). The monocytes shown in (D) are images of cells marked by corresponding numbers in the scatter plot. (E) The median monocyte cell size is significantly elevated in hospitalized COVID-19 patients (yellow, n = 17) compared to healthy blood donors (grey, n = 24) and recovered patients (green, n = 14). A significant increase is also observed in the standard deviation of cell size (F), median cell volume (G), and the standard deviation of cell volume (H) in COVID-19 patients compared to the healthy group. Statistical comparisons were done using Kruskal-Wallis test with Dunn’s posthoc test, * *p* < .05, ** *p* < .01, *** *p* < .001.

The differences in cell volume were also pronounced (*p* < .0001, *χ*^2^ = 27.7, ε^2^ = 0.51) with a 16.7% increase of median cell volume during COVID-19. The median volume of COVID-19 patient monocytes was 353.7 ± 55.8 µm^3^compared to 303.2 ± 12.0 µm^3^ for the healthy cohort and 304.9 ± 19.4 µm^3^for recovered patients (Figure 4 G). Assuming spherical shape, this would correspond to monocyte diameters of 8.8 µm for COVID-19 patients and 8.3 µm for healthy donors. Again, the standard deviation of cell volume was significantly higher for COVID-19 patients (*p* < .0001, *χ*^2^ = 18.5, ε^2^ = 0.34), as shown in Figure 4 H, due to the appearance of large, possibly highly phagocytic monocytes (19).

A morphological anomaly of COVID-19 patient monocytes has been observed by Zhang *et* al. (20). In that study, monocyte size was assessed indirectly using flow cytometry forward scatter (FSC). Unlike RT-DC, FSC measurement does not quantify absolute size changes (21). Still, the authors were able to observe a relative change of monocyte size and reported an increase in COVID-19 compared to healthy individuals, in line with our observation. In their study, the FSC-high population was more pronounced in patients requiring hospitalization and ICU admission.

No significant differences in deformation or Young’s modulus were found among the three studied groups (Supplementary figure 3), proving that the stiffness of monocytes remained unchanged during COVID-19.

### Altered physical phenotype signals neutrophil activation in COVID-19

Finally, RT-DC analysis provided evidence of significant changes with strong effect sizes in neutrophil cross-sectional area (*p* < .0001, *χ*^2^ = 23.0, ε^2^ = 0.43), volume (*p* < .0001, *χ*^2^ = 23.5, ε^2^ = 0.44) and deformation (*p* = .0013, *χ*^2^ = 13.3, ε^2^ = 0.25), Figure 5. During COVID-19, neutrophils were on average larger (68.7 ± 3.5 µm^2^ vs. healthy donors 63.5 ± 2.2 µm^2^, *p* < .0001), had higher volume (327.5 ± 27.2 µm^3^vs. healthy donors 292.0 ± 12.9 µm^3^, *p* < .0001) and were more deformed under the standard capillary flow conditions in RT-DC (0.059 ± 0.009 vs. healthy donors 0.051 ± 0.004, *p* = 0.002), Figure 5 F-H. The standard deviations of neutrophil cross-sectional area, volume and deformation were also significantly higher in the COVID-19 patients compared to healthy individuals (Supplementary figure 4). The joint increase of size and deformation of stimulated neutrophils *in vitro* and *in vivo* has been documented by RT-DC analysis previously (9, 22).

**Figure 5.**
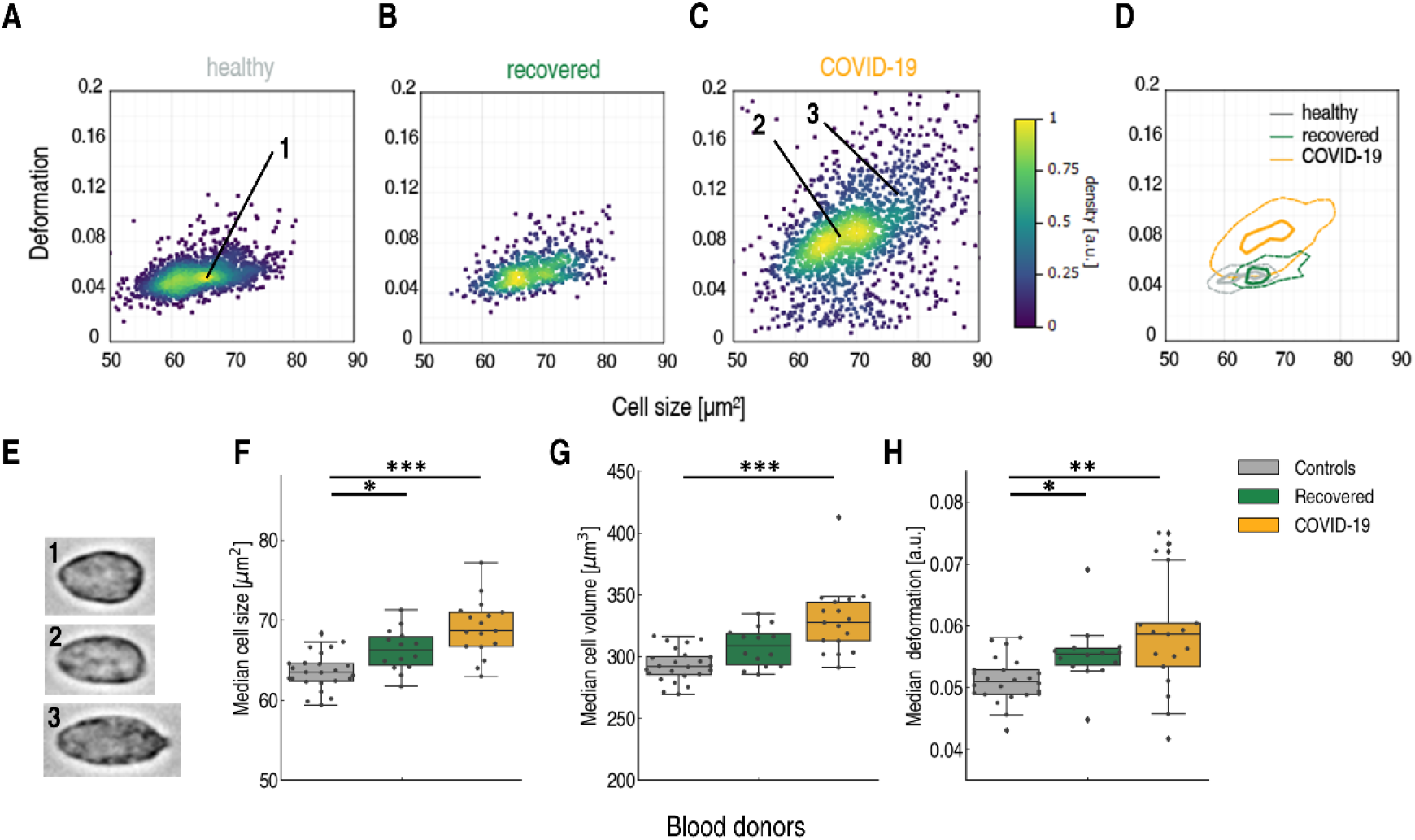
Altered physical phenotype of neutrophils in the peripheral blood of COVID-19 patients. Typical scatter plot of neutrophil deformation vs. cell size (cross-sectional area) of a healthy blood donor with no known viral infection (A) compared to a patient four months after undergoing COVID-19 (B) and a patient with COVID-19 in an intensive care unit (C). (D) Images of neutrophils marked by corresponding numbers in the scatter plots. (E) Kernel density estimate plots demonstrating the differences in cell size and deformation among the three donors (A-C). (F) The median cross-sectional area and (G) median cell volume of neutrophils of patients hospitalized with COVID-19 (yellow, n = 17) are significantly higher than that of the healthy blood donors (grey, n = 24) and of recovered patients approximately five months after undergoing COVID-19 (green, n = 14). (H) Neutrophils exhibit increased deformability in hospitalized COVID-19 patients compared to the healthy cohort. Statistical comparisons were done using Kruskal-Wallis test with Dunn’s posthoc test, * *p* < .05, ** *p* < .01, *** *p* < .001.

Thus, change in these parameters could serve as a proxy readout for neutrophil activation. We hypothesize that the changes of physical properties observed with RT-DC are linked to the strong activation and adoption of a low-density phenotype of neutrophils during COVID-19 (23).

The median Young’s modulus calculated from RT-DC data exhibited a weak decrease during COVID-19 (Figure 6 A), revealing that neutrophils were generally less stiff in COVID-19 patients. These changes of Young’s modulus were further confirmed by comparing data of three patients during and after COVID-19 (Figure 6 B), demonstrating a clear decrease of neutrophil stiffness during COVID-19. Interestingly, in recovered patients, neutrophil parameters (cross-sectional cell area 66.2 ± 2.5 µm^2^, volume 308.9 ± 15.3 µm^3^, deformation 0.055 ± 0.005) had not returned to the values of the healthy cohort (Figure 5 F-H). Carissimo *et* al. found that the neutrophil counts per defined volume of blood in recovered patients did not return to values of healthy individuals (24). Together with our findings, this suggests that COVID-19 infection leaves a lasting influence on the immune system.

**Figure 6.**
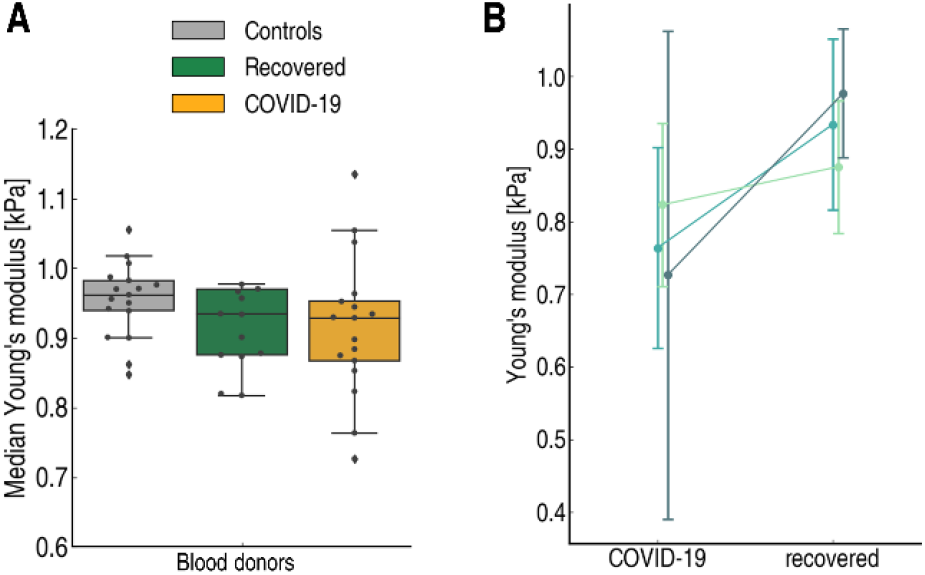
Decrease of neutrophil stiffness during COVID-19. (A) Young’s modulus of neutrophils of the three donor groups (healthy, recovered and COVID-19 patients). (B) Comparison of Young’s modulus of neutrophils in three patients measured at two time points: during COVID-19 and after recovery. Circle markers represent the median value, error bars represent standard deviation

In addition to neutrophils, we also examined a different group of granulocytes, eosinophils. Eosinophils are known to react to certain viral infections of the respiratory system *in vitro* and *in vivo*, including respiratory syncytial virus and influenza (25, 26). However, eosinophils did not show any changes of their physical phenotype during infection or in recovered state (Supplementary figure 5).

The above findings were reported as the medians and standard deviations of three mostly independent cohorts of blood donors. For three of the patients, we performed RT-DC measurements both during COVID-19 and after recovery, and could therefore directly examine the progression of blood cell parameters in a single individual. The comparison of various blood cell features of these three donors at the two studied time points can be found in Supplementary figure 6. The trends confirm what we described in the text above: severe COVID-19 is related to the presence of erythrocytes with distinct phenotype and lower deformation, larger monocytes, softer lymphocytes and neutrophils.

## Discussion

In our study, the physical parameters of blood cells including mechanical properties act as sensitive reporters of pathophysiological changes in COVID-19 patients compared to age-matched controls. The concept that cell morphology and mechanics are inherent markers of cell function has long been established (27, 28). Using RT-DC, we were able to monitor the physical properties of cells from whole blood without the need for tedious preparation or enrichment. We found alterations of erythrocytes and leukocyte subsets in COVID-19, which have the potential to be exploited as diagnostic markers. This paves the way for high-speed, label-free and cost-effective disease detection. We note that the relatively low number of COVID samples included in our study and the huge space of possible cellular responses to viral infection makes it a necessity to acquire many more RT-DC datasets to ensure the specificity of an observed pattern for a particular disease. Toepfner et al. provided initial evidence that the physical alterations of immune cells could distinguish a viral from a bacterial infection (9). In the future, RT-DC could be of high significance for fast distinction between viral and bacterial involvement. This would be an advantage in times when molecular diagnostic tests such as qPCR become inaccessible, such as during critical phases of a pandemic.

A key finding of our study was the altered physical phenotype of erythrocytes in COVID-19 patients. This is consistent with an erythrocyte proteomics report by Thomas et al., who identified structural protein damage and membrane lipid remodeling as potential causes of impaired oxygen delivery during COVID-19 (15). We found erythrocytes to be significantly more heterogeneous in shape and deformability under constant shear stress, compared to healthy controls. Moreover, the deformability of a significant subgroup of erythrocytes was dramatically reduced. The physical properties of erythrocytes are crucial for microcirculatory flow (29, 30) and as such, these changes could impair circulation and promote hypoxemia. The effect could persist in COVID-19 long haulers; we found that in recovered patients phenotype alterations were not as prominent, but still present. A different explanation for the persistence of less deformable erythrocytes in recovered patients could be that the cells already had a different physical phenotype before clinical onset. Altered mechanical properties of cells due to factors such as age could increase susceptibility to SARS-CoV-2 infection, as suggested by Uhler and Shivashankar (31).

While erythrocytes are present in much higher quantities (three orders of magnitude more frequent than leukocytes), also the numbers and physical features of leukocytes are crucial for proper blood flow (32). RT-DC provides direct access to relative leukocyte counts, their size and mechanical properties. In accordance with other studies (11, 33), we found neutrophilia and lymphopenia as well as increased neutrophil to lymphocyte ratio in COVID-19 patients. Importantly, we report on changes of size and mechanical properties of leukocyte subsets in COVID-19 samples. These changes might be key to understanding vessel occlusion and pulmonary embolism, as the interrogation of cell morphology and mechanics in former studies established the importance of these factors for cell circulation under physical (34), pathological (35, 36), and artificial (37) conditions.

Mechanical properties of cells can be directly related to the cytoskeleton (27, 28, 38, 39), an important supportive structure which also determines cellular function (40–43). Previously, using RT-DC we detected actin cytoskeletal rearrangements during rubella virus infection, which correlated with an altered cell shape and reduced migratory potential (44). Although it is known that viruses can use immune cells as vehicles to travel the body (45) and hijack the actin cytoskeleton (46), viral traces were absent in the blood of COVID-19 patients (47). However, the cytoskeleton may be affected by the infection indirectly *e*.*g*. through the involvement of cytoskeleton-dependent signaling (48). Hyperinflammation and cytokine storm syndrome are reported in COVID-19 cases with high levels of macrophage inflammatory protein 1-α, granulocyte-colony stimulating factor, interleukin (IL)-2, IL-7, interferon-γ inducible protein 10, monocyte chemoattractant protein 1, and tumour necrosis factor-α (49). Such cytokines were reported to induce cytoskeletal changes in myeloid cells and to interfere with their physical phenotypes during immune function (50–52).

Neutrophil degranulation and neutrophil extracellular trap formation were reported as a source of vascular occlusion, possibly leading to vascular damage and organ dysfunction in COVID-19 (23). This population of neutrophils is known to have a lower density, which could be logically linked with the elevated size and deformability observed with RT-DC. It is important to mention that neutrophils are short lived cells with an average lifespan of less than one day. Thus, neutrophil alterations observed in our recovered cohort were induced after the successful displacement of the virus by the immune system. This might be indicative for SARS-CoV-2 causing long-term immunological signals or even targeting bone marrow stem cells, as viral RNA was found *post-mortem* in patient bone marrow (53).

We are aware of certain limitations of our study, e.g. that it falls short of representative patient cohorts to compare mild with critical symptoms. Still, we were able to identify changes of blood cell physical phenotypes that did not return to the baseline healthy donor levels several months after SARS-CoV-2 infection, namely in erythrocytes and neutrophils. These alterations could be connected with long term symptoms of the recovered patients, of which 70% described chronic headache or neurological symptoms, 54% had concentration disorders and 62% circulatory problems like cold sweat and tachycardia. We hypothesize that the persisting changes of blood cell physical phenotypes could contribute to the impairment of circulation and oxygen delivery in coronavirus long haulers (17).

Taken together, label-free physical phenotyping of blood cells with real-time deformability cytometry provides a fast, sensitive and unbiased way to feel for functional changes in cells. As such, deformability cytometry data has the potential to be used as a biomarker of COVID-19 and potentially other infectious diseases. In the future, RT-DC could be part of the first line of defense against an unknown virus during a pandemic.

## Methods

### Peripheral blood collection

COVID-19 patients were hospitalized with a majority at intensive care unit (ICU) of the Department of Internal Medicine 1, Friedrich-Alexander-University Erlangen-Nürnberg, Germany (FAU) showing different severity levels at the time of blood sampling. COVID-19 venous blood samples (n = 17) were taken from hospitalized patients of the Department of Internal Medicine 1, FAU, between April and May 2020. Recovered patient venous blood samples (n = 14) were taken from patients of the Department of Internal Medicine 1, FAU, and COVID-19 patients in quarantine, in recovery from COVID-19 between four and eight months after release from the hospital or quarantine (median 7.1 ± 1.1 months). All patients had positive PCR tests for COVID-19. Control venous blood samples (healthy cohort) were taken from patients of the Department of Ophthalmology, FAU (patient information can be found in Supplementary table 1). Blood was drawn using a 20-gauge multifly needle into a sodium citrate S-monovette by vacuum aspiration with the tenets of the Declaration of Helsinki. Informed written consent was obtained from all participants. All experiments were performed according to the institutional guidelines and the ethical approval of the Ethical Committee of the University Medical Center of Erlangen (permits #193_13B and #174_20B and 295_20B). After blood collection, samples were used for clinical routine diagnostics and an aliquot was taken for RT-DC analysis within the standard storing time and conditions.

### Sample preparation

Prior to measurement, 50 µl of whole blood was gently mixed with 950 µl of measurement buffer (MB) composed of 0.6% (m/v) methyl cellulose dissolved in phosphate buffered saline (PBS; Figure 1 A), adjusted to a viscosity of 60 mPa.s at 24°C using a falling ball viscometer (Haake, Thermo Scientific).

### Real-time deformability cytometry

Real-time deformability cytometry (RT-DC) measurements were performed as described previously using an AcCellerator instrument (Zellmechanik Dresden GmbH) (9). The cell suspension was loaded into a 1 ml syringe, attached to a syringe pump (neMESYS, Cetoni GmbH) and connected by PEEK-tubing (IDEX Health & Science LLC) to a microfluidic chip made of PDMS bonded on cover glass. A second syringe with sheath fluid (pure measurement buffer) was connected to the chip, which consisted of two inlets (one for the sheath fluid and one for the sample) and one outlet connected by a channel constriction of 20 × 20 µm square cross-section, where the measurement was performed. The total flow rate was 0.06 µl/s, of which the sheath flow rate was 0.045 µl/s and the sample flow rate was 0.015 µl/s. To perform a measurement, the chip was mounted on the stage of an inverted high-speed microscope equipped with a CMOS camera. Measurement temperature was 23°C. Images were acquired at a frame rate of 1600 fps. Cells were detected in a region of interest of 250 × 80 pixels and morphological and mechanical parameters were acquired in real-time (Figure 1 B).

### Data analysis

Cell images were analyzed using ShapeOut software (54) and Python 3.7 using dclab library (55) For each patient, the six studied cell populations were hand-gated in the cell brightness-area parameter plot according to the procedure described in Toepfner *et* al. (9). The calculation of deformation, a measure of how much the cell shape deviates from circularity, and was obtained from the image using the projected area (*A*) and cell contour length calculated from the convex hull (*l*):

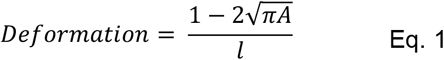

The calculation of the Young’s modulus was done using a look-up table derived from simulations based on the finite elements method (56) and the analytical solution (18). Cell volume was computed from the event contour under the assumption of rotational symmetry with a rotational axis parallel to the flow direction. The calculation is based on a full rotation of the upper and the lower halves of the contour, which are then averaged. Statistical analysis was done in Python 3.7 using Kruskal-Wallis *H*-test and post-hoc Dunn’s test with Bonferroni correction. In graphs, *P* values are represented by * *p* < .05, ** *p* < .01, *** *p* < .001. The effect size was estimated by the epsilon squared, ε^2^ (57), which was calculated from the *H*-test statistic as follows (58):

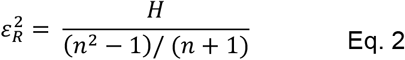

where *H* is the Kruskal-Wallis *H*-test statistic, n is the total number of observations, and the *ε*^*2*^coefficient assumes the value from 0 (indicating no relationship) to 1 (perfect relationship). The effect size was interpreted according to Rea *et* Parker (59), details are found in Supplementary table 2.

## Acknowledgements

The authors would like to thank Leonie Staats and Aylin Lindemann for technical assistance and Despina Soteriou, Michael Moritz, Charlotte Szewczykowski, and Folkert Horn for study support.

## Author contributions

M.K., J.G., and M.K. designed the project outline and carried out experiments, interpreted results, and wrote the initial manuscript. B.H., M.H. provided samples and patient information, interpreted and discussed results, and co-wrote the manuscript. J.H. and J.F. provided samples and patient information and co-wrote the manuscript.

## Conflict of interest

The authors declare no conflict of interest.

## Notes

### Competing Interest Statement

The authors have declared no competing interest.

