## Supplemental Figures and Tables for "Physical phenotype of blood cells is altered in COVID-19"

[1]Max Planck Institute for the Science of Light & Max-Planck-Zentrum für Physik und Medizin, Staudtstraße 2, 91058 Erlangen, Germany; [2]Department of Ophthalmology, Friedrich-Alexander-University Erlangen-Nürnberg, Schwabachanlage 6, 91054 Erlangen, Germany; [3]Department of Internal Medicine 1, University Medical Center Erlangen, Friedrich-Alexander-University Erlangen-Nürnberg, Erlangen, Germany; [4]Department of Internal Medicine 3, University Medical Center Erlangen, Friedrich-Alexander-University Erlangen-Nürnberg, Erlangen, Germany; [5]Deutsches Zentrum Immuntherapie (DZI), Erlangen, Germany; [6]Department of Physics, Friedrich-Alexander-University Erlangen-Nürnberg, Erlangen, Germany

### Supplementary Figures and Tables

**Supplementary table 1.** Patient characteristics, medical management and outcome of all donors included in this study.

| | All donors n=54 (100%) | | |
|---|---|---|---|
| | **Control n=24** | **Recovered n=14** | **COVID-19 n=17** |
| **Age (years): median (range)** | 62.5 ± 13.6 years (26-81) | 58.6 ± 12.4 (27-76) | 68·± 10.4 (41-87) |
| **Gender** | | | |
| male | 12 (50%) | 10 (71.4%) | 13 (76.5%) |
| female | 12 (50%) | 4 (18.6%) | 4 (23.5%) |
| **Primary virus identification (PCR airway)** | n.a. | 14 (100%) | 17 (100%) |
| **Complications and medical management** | | | |
| Oxygen supplementation | 0 | 0 | 17 (100%) |
| Mechanical ventilation | 0 | 0 | 13 (76.5%) |
| ECMO | 0 | 0 | 6 (35.3%) |
| Dialysis | 0 | 0 | 3 (17.6%) |
| Systemic Superinfection | 0 | 0 | 7 (41.1%) |
| Pulmonary embolism | 0 | 0 | 6 (35.3%) |
| **Drugs** | | | |
| Azithromycin | 0 | 0 | 3 (17.6%) |
| Hydroxychloroquine | 0 | 0 | 9 (52.9%) |
| Heparin prophylactic/therapeutic anticoagulation | 0 | 0 | 13 (76.5%) |
| **Outcome** | | | |
| Length of hospital stay (days) | 0 | 7 ± 2.4 (5-12) | 22.8 ± 14 (7-50) |
| Intensive care unit stay | 0 | 0 | 13 (76.5%) |
| Discharged | 0 | 14 (100%) | 9 (52.9%) |
| Further hospitalized | 0 | 0 | 0 |
| Death | 0 | 0 | 8 (47.1%) |

**Supplementary table 2.** Kruskal-Wallis *H*-statistics, *p*-values and effect sizes $\varepsilon^2$. The last three columns represent *p*-values from Dunn's posthoc tests conducted for the significant results of Kruskal-Wallis *H*-tests.

| A - healthy<br>B- recovered<br>C - COVID | | *p* (Kruskal-Wallis) | *H* (Kruskal-Wallis) | $\varepsilon^2$ (Kruskal-Wallis) | *p* AC (Dunn's) | *p* AB (Dunn's) | *p* BC (Dunn's) |
|---|---|---|---|---|---|---|---|
| **Erythrocytes** | | | | | | | |
| | **Median area** | 0.1162 | 4.30 | 0.080 | 0.4896 | 0.1503 | 1.0000 |
| | **SD area** | 0.0000 | 26.99 | 0.500 | 0.0000 | 0.2259 | 0.0112 |
| | **Median volume** | 0.2532 | 2.75 | 0.051 | 1.0000 | 0.2938 | 1.0000 |
| | **SD volume** | 0.0000 | 33.75 | 0.625 | 0.0000 | 0.9204 | 0.0002 |
| | **Median deformation** | 0.2180 | 3.05 | 0.056 | 0.4480 | 1.0000 | 0.3380 |
| | **SD deformation** | 0.0000 | 42.30 | 0.783 | 0.0000 | 0.0024 | 0.0340 |
| | **% of ery with def < 0.28** | 0.0000 | 25.83 | 0.478 | 0.0000 | 1.0000 | 0.0016 |
| **Neutrophils** | | | | | | | |
| | **Median area** | 0.0000 | 22.95 | 0.425 | 0.0000 | 0.2704 | 0.0260 |
| | **SD area** | 0.0001 | 18.86 | 0.349 | 0.0023 | 0.0001 | 0.6791 |
| | **Median volume** | 0.0000 | 23.53 | 0.436 | 0.0000 | 0.1319 | 0.0517 |
| | **SD volume** | 0.0001 | 19.78 | 0.366 | 0.0005 | 0.0002 | 1.0000 |
| | **Median deformation** | 0.0013 | 13.31 | 0.246 | 0.0021 | 1.0000 | 0.0319 |
| | **SD deformation** | 0.0059 | 10.28 | 0.190 | 0.0041 | 0.3772 | 0.5059 |
| | **Median Young's modulus** | 0.1698 | 3.55 | 0.066 | 0.1827 | 1.0000 | 0.8807 |
| **Lymphocytes** | | | | | | | |
| | **Median area** | 0.0499 | 6.00 | 0.111 | 0.1667 | 1.0000 | 0.0939 |
| | **SD area** | 0.0000 | 30.78 | 0.570 | 0.2270 | 0.0010 | 0.0000 |
| | **Median volume** | 0.0814 | 5.02 | 0.093 | 0.3403 | 1.0000 | 0.1134 |
| | **SD volume** | 0.0000 | 28.36 | 0.525 | 0.7403 | 0.0003 | 0.0000 |
| | **Median deformation** | 0.0132 | 8.66 | 0.160 | 0.0107 | 0.1945 | 1.0000 |
| | **SD deformation** | 0.0000 | 35.61 | 0.659 | 0.0043 | 0.0218 | 0.0000 |
| | **Median Young's modulus** | 0.0029 | 11.68 | 0.216 | 0.0029 | 0.0542 | 1.0000 |
| **Monocytes** | | | | | | | |
| | **Median area** | 0.0000 | 30.64 | 0.567 | 0.0000 | 0.0001 | 1.0000 |
| | **SD area** | 0.0011 | 13.65 | 0.253 | 0.0007 | 0.0892 | 0.7682 |
| | **Median volume** | 0.0000 | 27.71 | 0.513 | 0.0000 | 0.0001 | 1.0000 |
| | **SD volume** | 0.0001 | 18.48 | 0.342 | 0.0001 | 0.0075 | 1.0000 |
| | **Median deformation** | 0.7918 | 0.47 | 0.009 | 1.0000 | 1.0000 | 1.0000 |
| | **SD deformation** | 0.4949 | 1.41 | 0.026 | 0.7256 | 1.0000 | 1.0000 |
| | **Median Young's modulus** | 0.7763 | 0.51 | 0.009 | 1.0000 | 1.0000 | 1.0000 |
| **Eosinophils** | | | | | | | |
| | **Median area** | 0.1289 | 4.10 | 0.076 | 0.6099 | 0.1547 | 1.0000 |
| | **SD area** | 0.0126 | 8.7600 | 0.162 | 0.1245 | 0.0168 | 1.0000 |
| | **Median volume** | 0.2532 | 2.75 | 0.051 | 1.0000 | 0.2938 | 1.0000 |
| | **SD volume** | 0.0000 | 33.75 | 0.625 | 0.0000 | 0.9204 | 0.0002 |
| | **Median deformation** | 0.4143 | 1.76 | 0.033 | 1.0000 | 0.9095 | 0.5981 |
| | **SD deformation** | 0.5965 | 1.03 | 0.019 | 1.0000 | 1.0000 | 0.9290 |
| | **Median Young's modulus** | 0.9592 | 0.08 | 0.002 | 1.0000 | 1.0000 | 1.0000 |
| **% of WBC** | | | | | | | |
| | **% neutrophils** | 0.0105 | 9.10 | 0.169 | 0.0499 | 1.0000 | 0.0174 |
| | **% lymphocytes** | 0.0007 | 14.63 | 0.271 | 0.0006 | 0.2426 | 0.0328 |
| | **% monocytes** | 0.0151 | 8.38 | 0.155 | 0.0244 | 1.0000 | 0.0644 |
| | **% eosinophils** | 0.0010 | 13.73 | 0.254 | 0.1576 | 0.0007 | 0.1970 |
| | **NLR** | 0.0022 | 12.22 | 0.226 | 0.0031 | 0.7018 | 0.0252 |

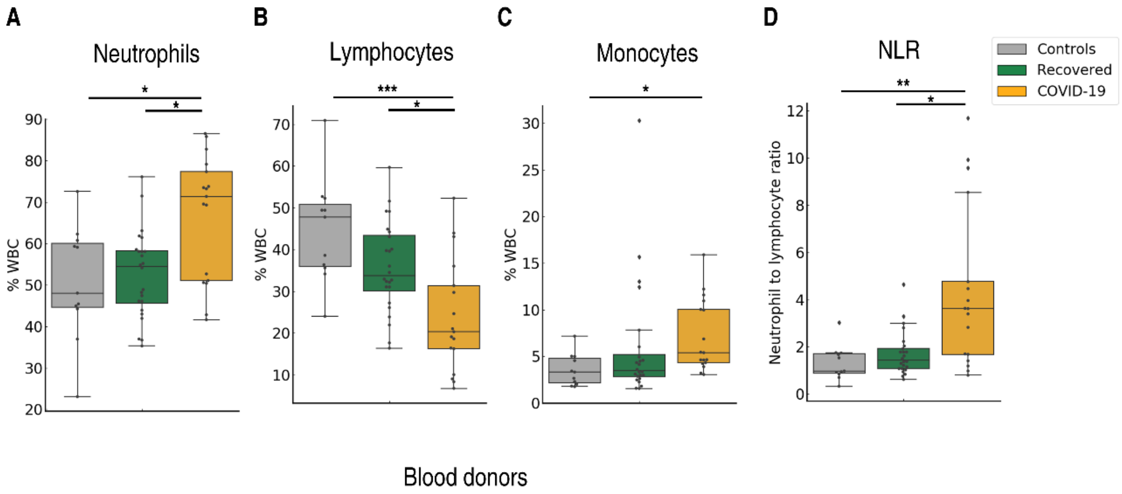

**Supplementary figure 1. Proportions of white blood cells calculated from real-time deformability cytometry (RT-DC) data.** The percentage of neutrophils (A), lymphocytes (B) and monocytes (C) in the total white blood cell count; a comparison of the control blood donor cohort (grey), recovered patients (green) and hospitalized COVID-19 patients (yellow). D) The neutrophil to lymphocyte ratio is significantly higher in hospitalized patients compared to the recovered and healthy donor cohorts, * $p < .05$, ** $p < .01$, *** $p < .001$.

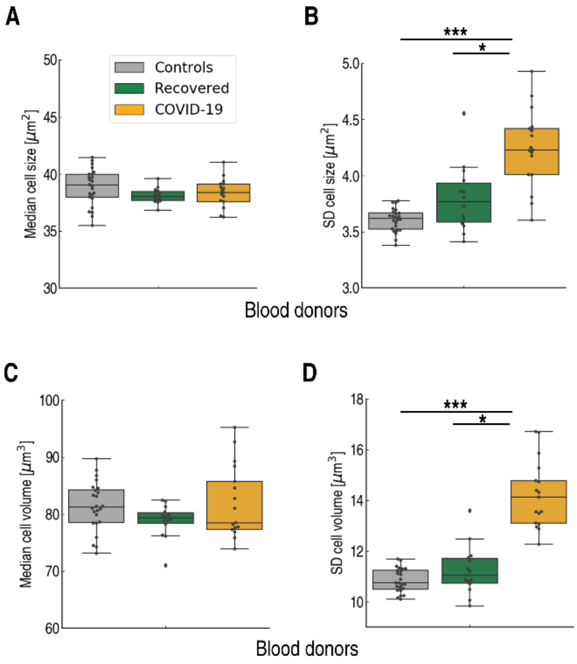

**Supplementary figure 2. Cell size of erythrocytes of COVID-19 patients compared to controls.** The graphs compare the control group of blood donors (n = 24), recovered patients (n = 14), and hospitalized COVID-19 patients (n = 17). While the median of cell size is similar in the three groups (A), the standard deviation of cell size is significantly increased in COVID-19 patients (B). In similar fashion, the volume of erythrocytes remains unchanged (C) but the standard deviation of cell volume is significantly higher during COVID-19 (D). Statistical comparisons were done using Kruskal-Wallis test with Dunn's posthoc test, * $p < .05$, ** $p < .01$, *** $p < .001$.

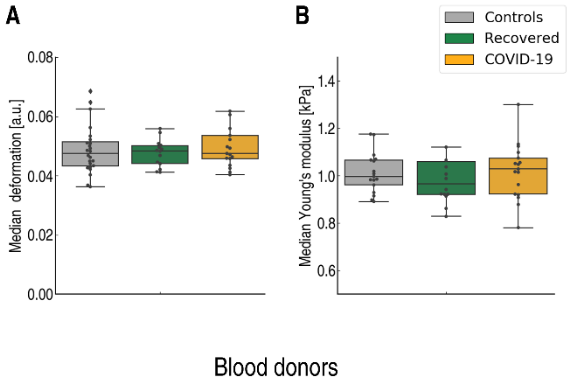

**Supplementary figure 3. A) Deformability and B) Young's modulus of monocytes of COVID-19 patients compared to controls.** The graphs compare the control group of blood donors (n = 24), recovered patients (n = 14), and hospitalized COVID-19 patients (n = 17). No statistically signifiant differences were found. Statistical comparisons were done using Kruskal-Wallis test with Dunn's posthoc test, * $p < .05$, ** $p < .01$, *** $p < .001$.

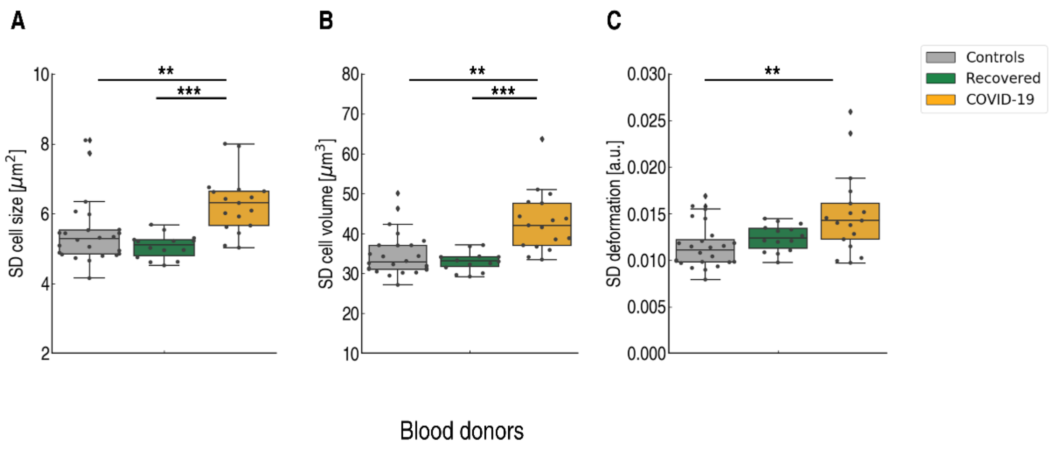

**Supplementary figure 4. Standard deviations of cross-sectional cell size (A), volume (B) and deformation (C) of neutrophils.** The graphs compare the control group of blood donors (n = 24), recovered patients (n = 14), and hospitalized COVID-19 patients (n = 17). The standard deviations of the parameters increase during COVID-19, differences between the healthy cohort and hospitalized COVID-19 patients were statistically significant. Statistical comparisons were done using Kruskal-Wallis test with Dunn's posthoc test, * $p < .05$, ** $p < .01$, *** $p < .001$.

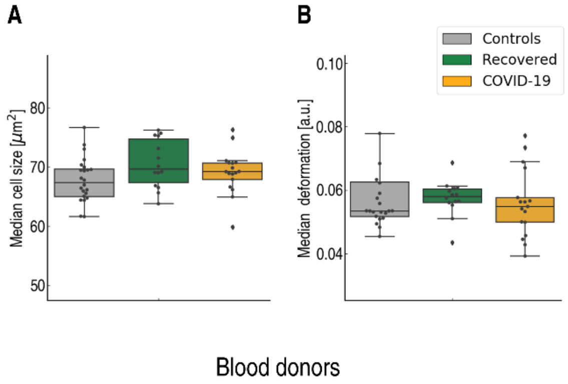

**Supplementary figure 5. Median values of cell size and deformation of eosinophils of COVID-19 hospitalized patients compared to recovered patients and healthy donors.** Differences between the three groups were not significant. Statistical comparisons were done using Kruskal-Wallis test with Dunn's posthoc test, * $p < .05$, ** $p < .01$, *** $p < .001$.

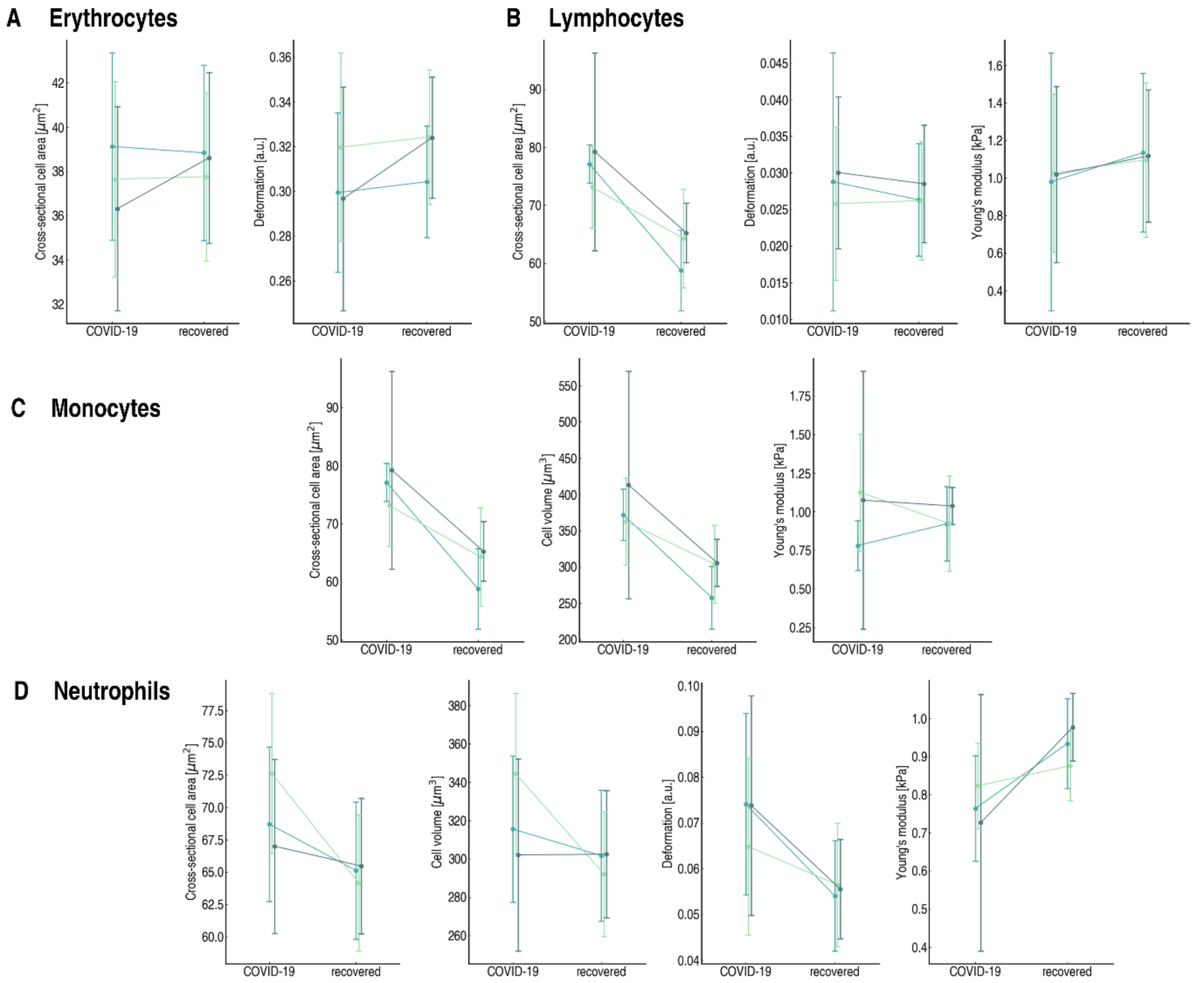

**Supplementary figure 6. Comparison of the blood cell parameters of three different patients measured at two time points: during COVID-19 and after recovery.** (A) Erythrocytes, (B) lymphocytes, (C) monocytes, (D) neutrophils. Presented are the most interesting blood cell parameters discussed in the manuscript. Circle markers represent the median value, error bars represent the standard deviation.
